# Tetra hydroxy ethyl disulfate disodium: a novel non-resistant antibacterial chelating agent

**DOI:** 10.1101/2024.06.04.597394

**Authors:** KA Audah, GE Jogia, D Kustaryono, J Amsyir, SA Amalda, J Darmadi, S Himawan, S Saidin, BA Tejo, C Budiman, S K Singh, J Thekkiniath

## Abstract

Discovery of novel antibacterial drugs is urgently required as common antibiotics have numerous side effects and are becoming less effective due to antibiotic resistance occurrences. Developing a novel class of antibacterial compounds without inducing the prevalence of resistance when used repeatedly against bacterial pathogens is presumably one of the best approaches to overcome this problem. Tetra hydroxy ethyl disulfate disodium (THES) is a novel antibacterial agent, capable of disrupting cell activity without any need for drug influx. In this study, the molecular interaction between THES, Gram-positive, and Gram-negative bacteria was investigated. It was suggested that THES possessed the desirable characteristics of a non-resistant bacterial agent with bactericidal properties. Additionally, the non-resistant characteristics of THES were validated as a novel antibacterial agent. The unique feature of THES was the property of chelation with its strong binding ability to target ligands. This binding played a major role in bacterial peptidoglycan porosity enlargement, resulting in lysis and cell death. The bacterial cell membrane porosity enlargement was confirmed by Scanning Electron Microscopy, Transmission Electron Microscopy, as well as Atomic Force Microscopy. These findings were further supported by the results from *in silico* analysis which showed that THES formed favorable interactions with peptidoglycan with a binding affinity of −4.7 kcal/mol. The minimum inhibitory concentration value against multidrug-resistant (MDR) bacteria was notably low, between 0.05 - 0.1%. Furthermore, the analysis of resistance over a seven-month course showed that *Staphylococcus aureus* did not develop resistant characteristics to repeated treatment to THES treatment. This study has also shown that bactericidal characteristics can be easily manipulated and modified for more preferable antibacterial activities, whilst yielding desirable pharmacokinetic/pharmacodynamics (PK/PD) properties. There are several advantages of THES as compared to common antibiotics, including its flexibility in the killing capacity of different microbial strains. The simplicity, efficacy, and non-resistant properties of THES make it a promising drug candidate or compound to combat the outbreak of MDR bacteria in the future.

## Introduction

Negative human reactions to bacterial infections by the excessive use of antibiotics [1–4], followed by the application of high standards in hygiene and sanitation have led to bacterial mutation and survival which induce bacterial resistance to commonly used antibiotics [3]. Widespread and inappropriate overuse of antibiotics, including those outside of human medicine, such as in agriculture [5] and aquaculture, have contributed to the rapid evolution and spread of antibiotic-resistant bacterial pathogens that have now threatened our ability to control bacterial infections [6,7]. Inevitably, with the increasing use of antibiotics, more antimicrobial resistance (AMR) traits are observed in bacteria. Despite the reduction of inappropriate antibiotic use has slowed down this problem, it does not halt the growing number of AMR bacteria and microbial strains. Existing antibiotics continue to lose their effectiveness over time, thus infected patients will continuously require new drugs and therapies. The growing frequency of resistant bacterial outbreaks has prompted significant research into the development of new antibiotics with novel targets [8–10].

A new strategy for modifying existing antibiotics and antibacterials is required to confer improved activity, less sensitivity to resistance mechanisms, and less toxicity [11–13]. One interesting bactericidal activity that bypasses the resistance development is by influencing bacterial morphology and physiology from outside the bacteria, such as alcohol and its derivatives by dehydrating the bacterial cytoplasm [14–16]. However, the usage of alcohol as a medicine is restrictive due to its irritability towards human tissues [17,18]. Since it would necessitate a change in the fundamental chemical properties of biological molecules, human tissue immunity to alcohol has not been observed to evolve [17]. Therefore, antibacterial agents possessing a specific killing mechanism that can work from outside the bacteria need to be relatively safe, so as not to injure surrounding non-target tissues and cells.

The most economically valuable antibacterial agent should be easily synthesized, modified readily, taken orally, has broad bioavailability, low toxicity, and can be prescribed for a wide spectrum of bacterial pathogens [19,20]. Novel classes of antibacterial drugs should be tested for antimicrobial activity against non-attenuated bacterial strains of the initial test species, as well as against other relevant pathogenic microorganisms such as fungal specimens and some antibiotic-resistant bacterial strains [21]. The goal of the *in vitro* study is to identify a drug with low minimum inhibition concentration (MIC), preferably against more than one pathogen [20]. An *in vivo* study to confirm safe application against a mammalian model should also be performed [22]. Finally, resistance studies are necessary to determine the frequency or rate of spontaneous resistance development to the progressed compounds as a function of drug exposure and the relative fitness of resistant mutants by using strains of the target profile pathogens [23, 24]. In this study, a newly synthesized compound, a patented product, tetra hydroxy ethyl disulfate disodium (THES) was proposed as a novel nonresistant antibacterial chelating agent. The structure of THES as depicted in Fig. 1 was elucidated by the FTIR experiment.

**Fig. 1.**
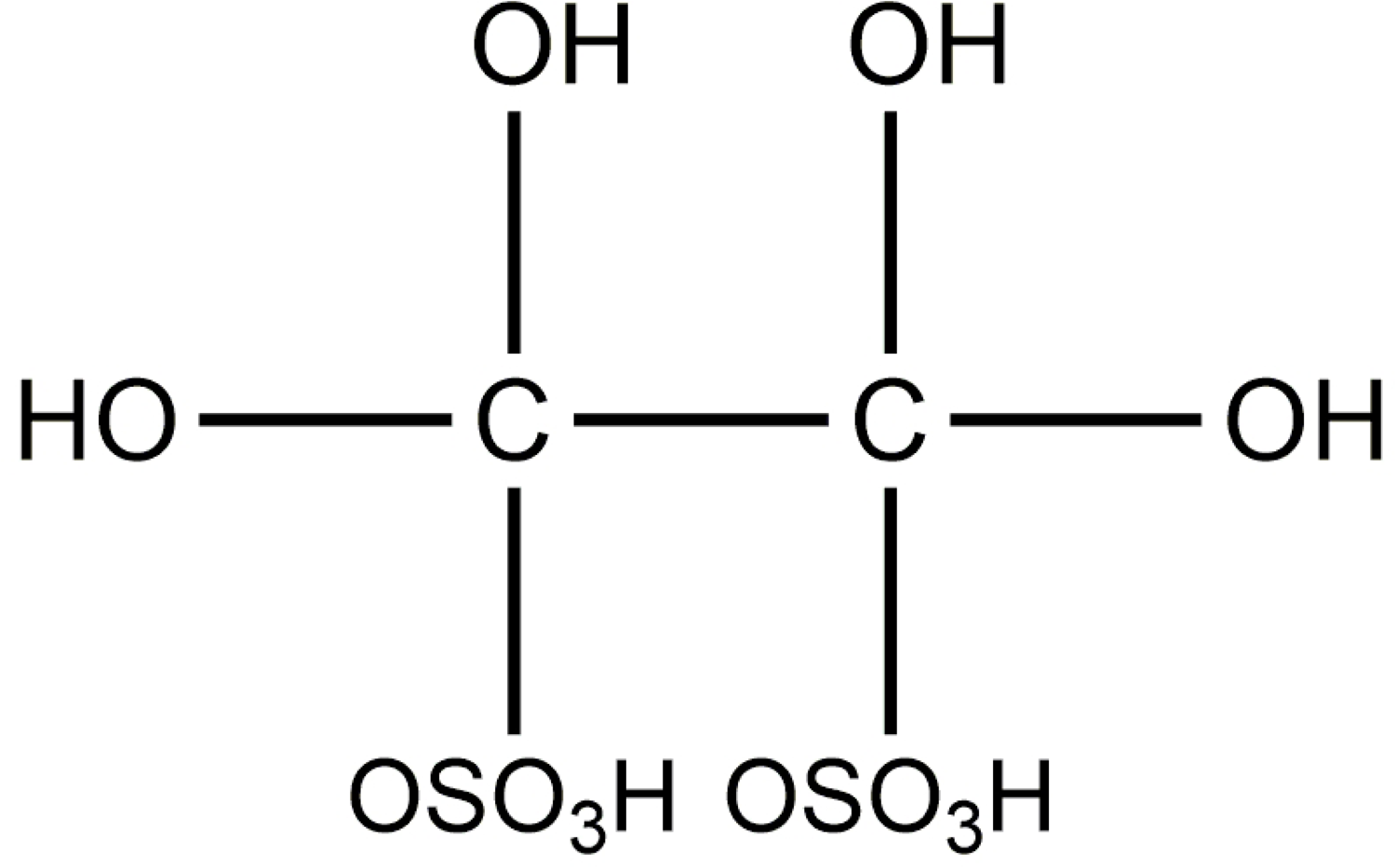
THES chemical structure

THES hypothetically works on the peptidoglycan cell membrane of bacteria by loosening or completely cutting the cross-linking of carboxyl amino bonds between peptides in protein or hydroxy-hydroxy bonds between glycan in the peptidoglycan structure. THES’s unique property as a sulfate chelating agent, which is contributed by its extremely strong binding property with the target ligands, is the primary cause of increased peptidoglycan membrane porosity in bacteria, resulting in lysis and eventually cell death.

The study for THES activity was conducted through *in silico* and *in vitro* tests as a complex process for bringing THES to be an acceptable antibacterial agent. However, the risk and cost of developing a new class are significantly higher than those of an analogue [25]. The study included *in silico* binding study between THES and peptidoglycan as a major bacterial cell membrane component. *In vitro* activity against multidrug-resistant (MDR) bacteria was performed by measuring MIC value as well as a binding study between THES and peptidoglycan by using the Isothermal Titration Calorimetry (ITC). The bacteria peptidoglycan porosity enlargement was confirmed by Scanning Electron Microscopy (SEM) and Transmission Electron Microscopy (TEM) imaging. The resistance test was finally conducted on *Staphylococcus aureus* where the bacteria was treated with THES for seven months.

## Materials and Methods

### Bacterial strains and fungal samples

*Salmonella typhi*, *Staphylococcus aureus* ATCC 6538, and *Escherichia coli* ATCC 9637 were obtained from Microbiology laboratory, Pharmacy School of Bandung Institute of Technology. *Bacillus subtilis* ATCC 6633, *Staphylococcus epidermidis* ATCC 12228, *Corynebacterium diptheriae*, *Propionibacterium acne*, *Acinetobacter anitratus*, *Pseudomonas aeruginosa* ATCC 27853, *E. coli* ATCC 25922, *Aspergillus niger*, and *Candida albicans* ATCC 10231 were obtained from Cell Culture Collection, University of Indonesia. MDR strain bacteria Vancomycin-resistant *Enterococcus faecalis*, Carbapenem-resistant *Klebsiella pneumonia*, and β-lactam-resistant (NDM-1 positive strain) *E. coli* were obtained from Bioscience Laboratory Montana.

### Chemicals and reagents

Tetra hydroxy ethyl disulfate disodium (THES) is a patented product with patent numbers: WO2014097284A1, US20150315135A1 (PT. Novis Natura Navita, Tangerang, Indonesia). Other reagents or chemicals include nutrient broth and nutrient agar (Difco), Tryptic Soy Agar, ethanol 70%, sterilized aquadest, glutaraldehyde, bovine serum albumin (BSA), phosphate buffer (pH 7), 0.9% sodium chloride (NaCl), LR white resin, toluidine blue, sodium citrate trihydrate (C_6_H_11_Na_3_O_10_), 0.1 N sodium hydroxide (NaOH), uranyl acetate (C_4_H_6_O_6_U), lead nitrate (Pb(NO_₃_)_₂_), pyrogen-free sterile saline, rubbing alcohol, phenol, peptidoglycan, 20 mM MES (pH 6.8), 100 mM NaCl, 2 mM CaCl_2_ buffer, and KBr.

## Methods

### 1. FTIR Spectroscopy of Synthesized THES

Synthesis of THES is part of the patented product, hence it is not written in this manuscript. The resulting THES compounds were then dried at 60°C and 100°C using KBr. The dried THES compounds were read using FTIR– Bruker Alpha. Furthermore, the resulting chromatograms were compared with the IR table. The FTIR-Bruker Alpha tool showed the THES functional group at a wavelength between 500-4000 cm^-1^.

### 2. In silico study

#### A) Software used

Avogadro (Avogadro 1.97.0 documentation, Avogadro Teams) was used to construct the three-dimensional structure of the ligand [26]. BIOVIA Discovery Studio Visualizer (Dassault Systèmes Biovia, Dassault Aviation, French) was utilized to visualize and modify the receptor and the ligand structure, as well as post-docking interaction analysis. OpenBabel is a molecular format conversion program used to convert various ligand formats [26]. AutoDock Vina was the main docking software used in this work [27,28]. AutoDock Tools (The Scripps Research Institute, La Jolla, USA) was used for the preparation of the .pdbqt files of protein and ligands.

#### B) Peptidoglycan preparation for molecular docking

The nuclear magnetic resonance (NMR) structures of peptidoglycan in complex with *B. subtilis* L, D-transpeptidase were downloaded from the RCSB Protein Data Bank (PDB ID: 2MTZ) (http://www.rcsb.org/). The peptidoglycan structure was set for further molecular docking by eliminating the transpeptidase using BIOVIA Discovery Studio Visualizer.

#### C) Preparation of THES ligand for molecular docking

The three-dimensional structure of THES was prepared using Avogadro software. The structure was optimized using the MMFF94 force field with the steepest descent algorithm for 500 steps with the convergence criteria of 10^-7^.

#### D) Molecular docking

AutoDock Tool was used to prepare the .pdbqt input file for peptidoglycan and to determine the size and the center of the gridbox. Polar hydrogen atoms and Kollman charges were set for peptidoglycan. The center of the gridbox was set to −7.856 × −1.953 × 1.828 in the dimensions of x, y, and z, respectively using 1.000 Å spacing and a box size of 40 x 40 x 40. Molecular docking was performed using AutoDock Vina. The predicted binding affinity (kcal/mol), which indicates how strongly THES binds to peptidoglycan, was calculated based on the scoring function used in AutoDock Vina. BIOVIA Discovery Studio Visualizer was used to analyze the docking results.

### 3. *In vitro* activity of THES

*In vitro* activity of THES was conducted at Bioscience Laboratory Montana, USA. The following procedures were used to conduct *in vitro* activity. The THES solutions at concentrations of 100 and 1000 µg/L were made by diluting the stock solution (10,000 mg/L). There were two rows of 12 sterile 7.5 x 1.37 cm screw-capped tubes. One row was inoculated by one drop of an overnight broth culture of the test organism which has been diluted to 1 in 1000 in a nutrient broth and the second row was inoculated with the control organism of known sensitivity similarly diluted. The size of the inoculum has a significant impact on the test’s outcome. 10^6^ CFU/ml should be presented in the test mixture. If there was not enough growth, the undiluted version would be used as a replacement. The tubes were incubated at 37°C for 18 hours. The 2 ml broth with the organism was inoculated and stored at 4°C overnight to serve as a standard for determining complete inhibition.

### 4. TEM and SEM

The TEM examination was carried out at the Pharmacy School of the Bandung Institute of Technology. *S*. *aureus* and *E. coli* were cultured in nutrient broth for 24 hours at 37^°^C. 1.5 ml culture was transferred to microcentrifuge vials, and 0.5 ml of 0.6, 5, and 10% THES was added and incubated for four hours. The samples were centrifuged and rinsed with 0.9% NaCl and distilled water, respectively. The pellets were fixated in 2% formaldehyde and 0.5% glutaraldehyde for two hours at 4^°^C. The pellets were rinsed thrice with phosphate buffer pH 7, pre-embedded in bovine serum albumin (BSA) 1/5, and were then allowed to harden at room temperature. The microbes were dehydrated with alcohol, then infiltrated with resin LR White and polymerized for 48 hours at 60^°^C to form hard-capsule-like products. The infiltrated resins were sectioned using an automatic ultra-microtome. Semi-thin sections were prepared, stained with toluidine blue then examined under a microscope to obtain the right section. After obtaining the exact section, ultra-thin sections were made and contrasted with uranyl acetate and lead citrate. The sections and samples which have been dried were examined under TEM.

The SEM observation was conducted at the Research Center for Physics, Indonesian Institute of Sciences. *S*. *aureus* and *E. coli* were cultured in Mueller Hinton broth for 24 hours at 37^°^C. 1.5 ml culture was transferred to microcentrifuge vials, 0.5 ml 0.1% THES was added, and the vials were left to stand for 30 minutes. The samples were centrifuged, with supernatant discarded and the pellets were rinsed with distilled water. The pellets were fixed in 4% glutaraldehyde at 4^°^C and then were centrifuged after 30 minutes, with the supernatant discarded. All pellets were resuspended in 0.1 M phosphate buffer for 20 minutes and centrifuged to wash the pellets. The pellets were resuspended again in 0.1 M phosphate buffer containing 1% osmium tetroxide. After one hour, the suspensions were centrifuged and supernatants were discarded. The pellets were resuspended in distilled water and washed again through centrifugation. The samples were then dehydrated using an ethanol series (35, 50, 75, and 95%) and HDMS as follows, centrifuging and discarding the supernatant after each change. The supernatants from the second HDMS were discarded, and the vials containing the cells were left overnight in a desiccator to air dry at room temperature. After that, the dried cells were mounted on the SEM stub. The samples were sputtered with gold powders and examined under the SEM.

### 5. Observation of THES effect on bacterial membrane morphology using AFM

The use of AFM as a measuring technique is well established and discussed in several published literature [29–32]. It has been used to study the morphology and topography of cells, including structural and functional cell surface, biophysical, surface adhesion, and surface roughness [33–35].

Before imaging samples, the fixation of cells using attaching agents was often used to prevent artifacts from external movement and noise from the environment and eliminate the mobility of viable biological samples, such as cells, bacteria, fungi, and other microorganisms. Poly-L-lysine, a non-specific attachment factor cell, was used as an attaching agent in this study. Poly-L-lysine was chosen because of its ability to keep biological samples viable during the imaging process, but it may flatten or spread to such an extent that it can be ignored for observational purposes [36, 37].

The principle of its attachment property lies in its electrostatic interaction between the negatively charged cell membrane and the polymer cationic amino acid which contains approximately one hydro-bromide (HBr) molecule per lysine residue. The HBr group will interfere with hydrogen bonding between amino and either the carboxyl, nitrogen, or oxygen-containing moieties. Fixation of cells using poly-L-lysine with higher molecular weight (150,000– 300,000) is preferable to provide more attachment per molecule. The polycationic molecules will adsorb strongly to solid surfaces, leaving cationic sites that combine with the anionic sites on cell surfaces.

The AFM images obtained in this study provided geometrical and topographical information, allowing for quick analysis of objects and organized measurements of individual samples. By defining lower and upper color threshold levels ranging from 59.5 nm (black) to 296 nm (black), presenting data ranging from 90.6 nm to 249.0 nm for *E. coli* specimen; 14 nm (black) to 549.7 nm (white), presenting data ranging from 61.1 nm to 555.3 nm for *S. aureus* specimen (control); and 14.60 nm (black) to 802.8 nm (white), presenting data ranging from 61.1 nm to 910 nm for *S. aureus* specimens exposed to the THES.

To reduce the possibility of false data presentation due to the tilt from the sample support, the lower sample as in the background for the poly-L-lysine in the data was removed using the ‘Plane Fit function. For the control sample, the images were captured in air mode immediately, and for the treatment sample, 35 minutes after the THES-treated sample was incubated. Samples were observed at a scan area of 3 x 3 μm^2^, allowing individual assessment of the sample. The THES-treated sample was conditioned with 0.5% THES for 35 minutes on a poly-L-lysine-coated slide. The scan area of the sample containing *E. coli* treated with THES was narrowed to 2 x 2 μm^2^ since the scanning area of 3 x 3 μm^2^ was considerably too wide as a scan area compared to the focus area to be observed.

### 6. Peptidoglycan binding test of THES

The Peptidoglycan binding test of THES was conducted by using Isothermal Titration Calorimetry. ITC experiments were performed using a titration microcalorimeter called MicroCal ITC200 (MicroCal Inc., Northampton, MA). Peptidoglycan was dialyzed simultaneously in 20 mM MES, pH 6.8, 100 mM NaCl, and 2 mM CaCl_2_ buffer. The peptidoglycan concentration in the sample cell was 30 μM in a 0.2 ml total volume, and 1.5μl of the compound was sequentially injected into the sample cell for 25 injections with a ligand-protein concentration of 380 µM. This experiment was carried out on the sample cell and reference cell at 25°C. The heat of the reaction was obtained by integrating the peak after each injection of ligand-protein using Origin software provided by the manufacturer. In this experiment, BSA was used as a negative control with the same treatment as peptidoglycan. *K_d_*value as the indicator of binding strength was calculated according to the following formula [38]:

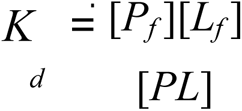

### 7. Resistance and Antiseptic Test

THES resistance test was performed using the Rideal-Walker phenol coefficient experiment for seven-month periods [39]. Pure *S. aureus* colony culture was suspended in nutrient broth and incubated for 18-24 hours at 37^°^C. At 530 nm, an intermediate *S. aureus* suspension was created in 25% NB. 26% THES solution was used to make different concentrations of THES in water (0.5-10 mg/mL or dilution ratio 1:200-1:10) and were further placed in sterile reaction tubes. The *S. aureus* bacterial suspension was transferred into each THES test tube reaction solution using an inoculation loop and the tubes were further vortexed. Immediately after the short vortex, the contact time between *S. aureus* and THES solutions was 2, 5, and 10 minutes. Phenol solutions with concentrations 1.67-0.5 mg/mL or dilution ratio 1:60-1:200 were also used as positive controls. An inoculation loop was then used to collect the treated bacterial samples after contact time 2, 5, and 10 minutes into duplicates of 5 ml sterile medium NB to be further incubated at 37°C for 48 hours. After each month, bacterial suspension was cultured in NB that was added with a concentration of 0.3% THES MIC. The phenol coefficient of THES each month was calculated by dividing the highest achievable dilution ratio of THES over the dilution ratio of phenol that resulted in bacterial death after 10 minutes of contact time yet did not result in death within 5 minutes.

## Results

### 1. FTIR analysis of THES synthesis

On the dried THES at 60°C (Fig. 2), the spectral peak of 853.74 cm^-1^ was identified to correspond to the C–C bond. The next peaks to be identified were at 1040.74 - 1232.53 cm^-1^ (C–O), 1307.82 cm^-1^ (sulfonate), and 3497.34 cm^-1^ (O–H). Meanwhile, the THES sample dried at 100°C (Fig. 3) was found to have peaks at 843.40 - 995.10 cm^-1^ which corresponds to the C–C bond, then 1070.85 - 1234.38 cm^-1^ (C-O), as well as 1333.45 cm^-1^ which was also a sulfonate functional group peak.

**Fig. 2.**
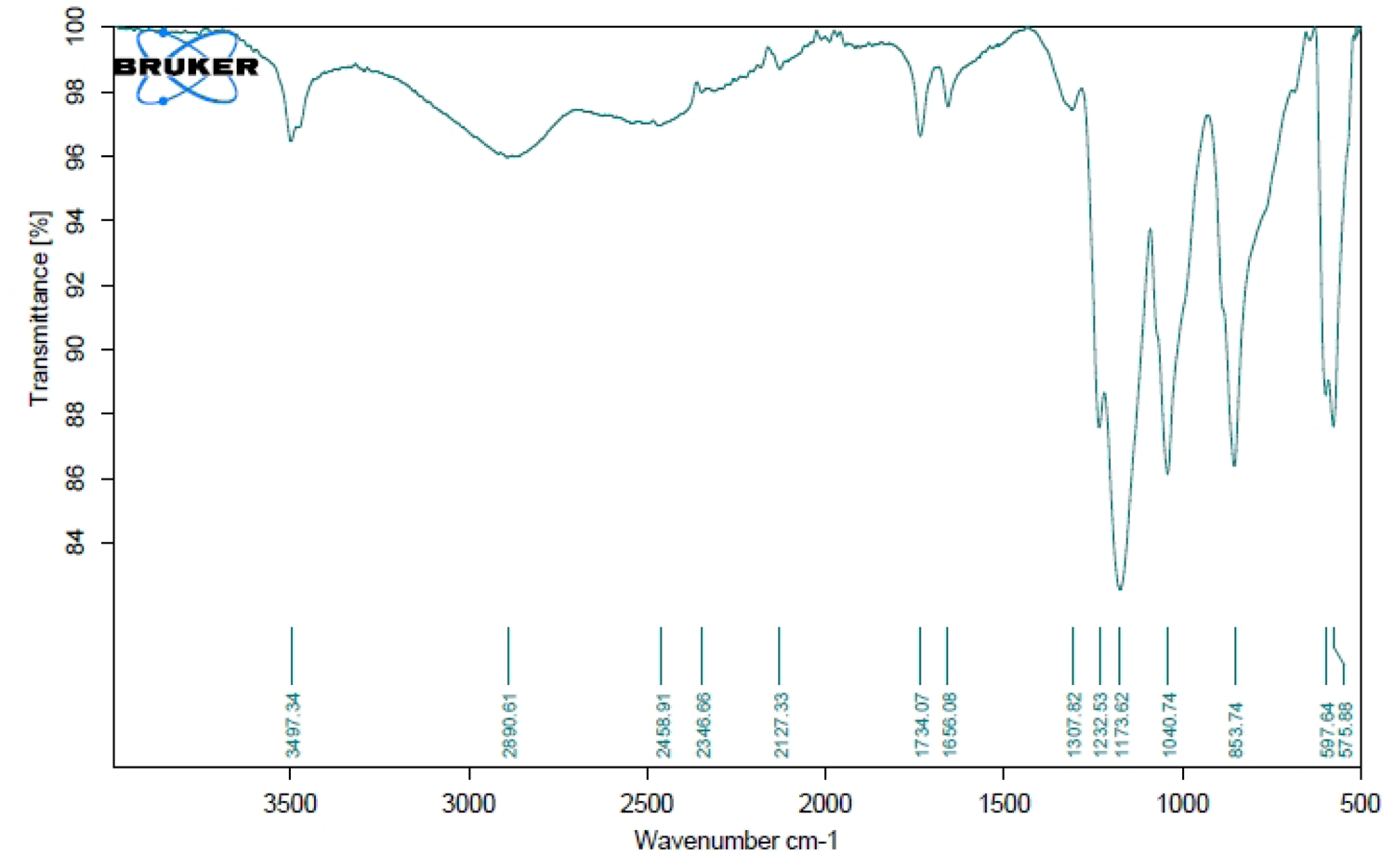
THES FTIR Chromatogram with Drying at 60°C.

**Fig. 3.**
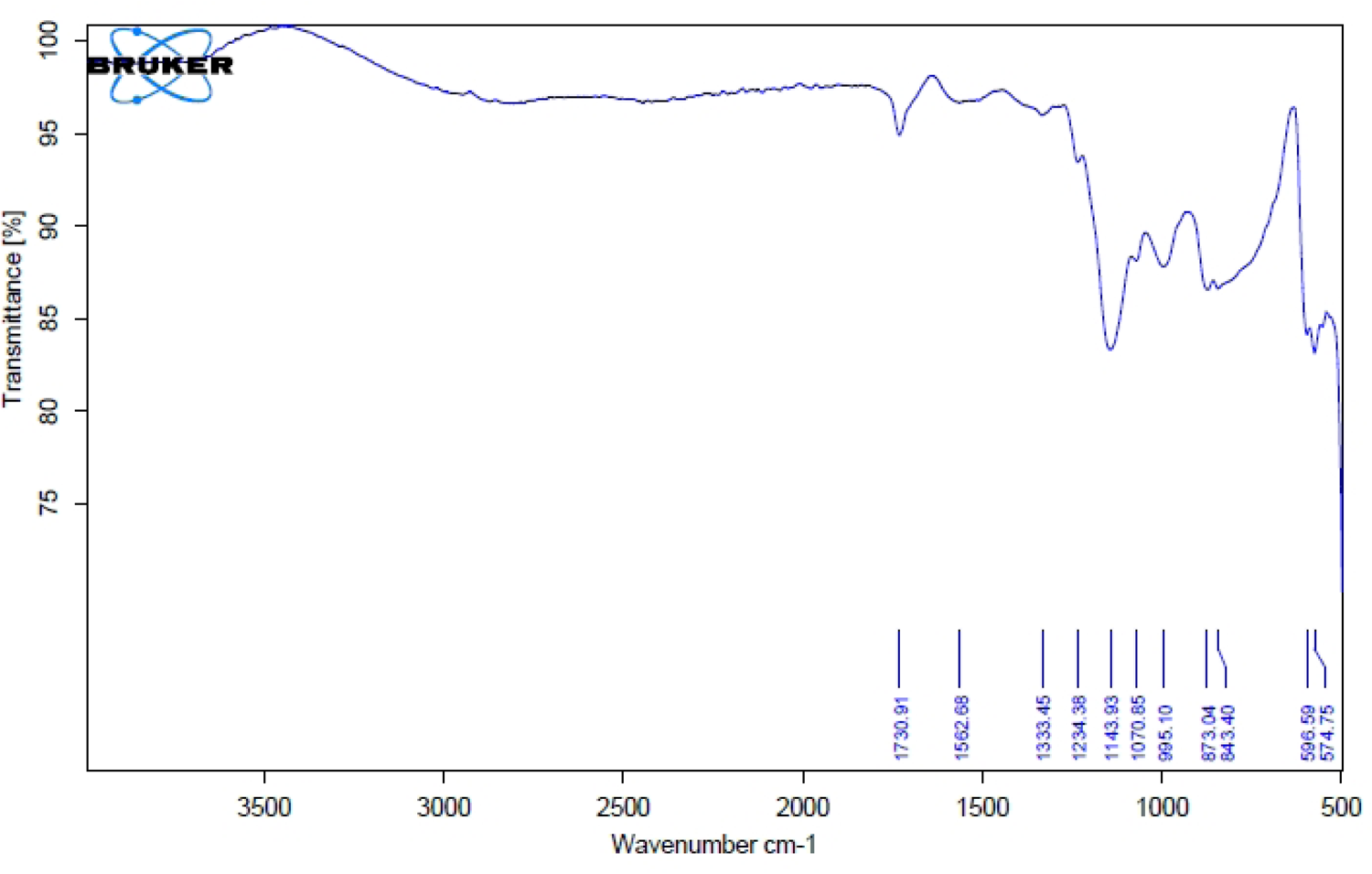
THES FTIR Chromatogram with Drying at 100°C.

### 2. Interaction between THES and peptidoglycan was determined using molecular docking

Our molecular docking result showed that THES forms favorable interaction with peptidoglycan with a binding affinity of −4.7 kcal/mol. THES was stabilized by multiple hydrogen bonds between different residues of peptidoglycan (Fig. 4). The O6 oxygen atom of residue NAG2 at chain H formed the most hydrogen bonds with THES, i.e. atom O6 in the carbonyl group, atoms O3 and O9 in the hydroxy groups, and atom H11 in the hydroxy group. Residue FGA34 at chain G of peptidoglycan formed two hydrogen bonds with atoms H31 and H41 of the hydroxy groups. The N6 nitrogen atom of residue API29 at chain F formed a hydrogen bond with atom O6 at the carbonyl group of THES. The O3 oxygen atom of residue NAM1 at chain H formed a hydrogen bond with atom H91 of the hydroxy group.

**Fig. 4.**
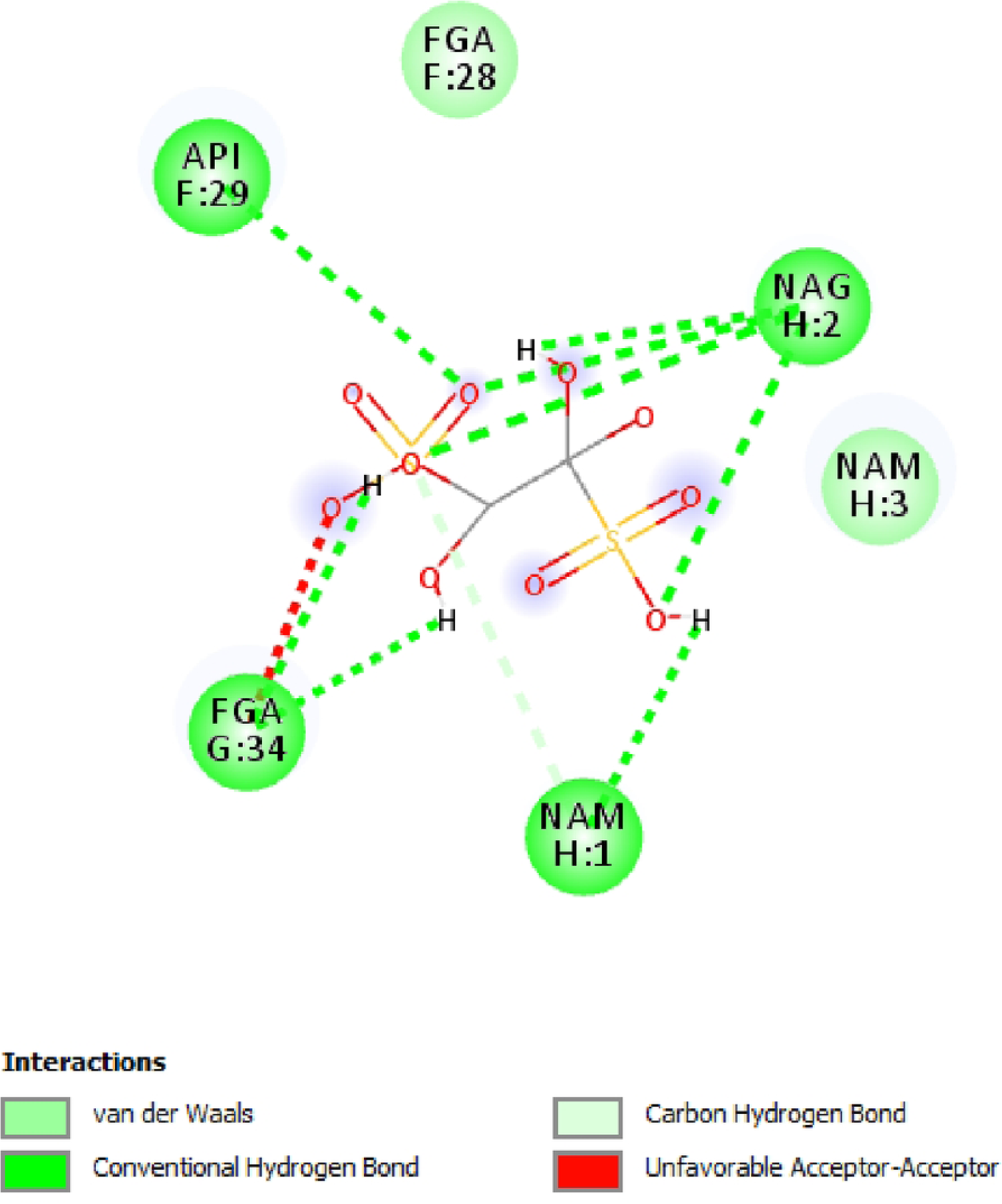
Binding conformation of THES and its interactions with the surrounding peptidoglycan residues.

### 3. *In-vitro* Activity Test

THES activity was tested *in vitro* against a variety of microorganisms, including both Gram-negative and Gram-positive bacteria. The test included Vancomycin-resistant *E. faecalis*, carbapenem-resistant *K. pneumonia*, and β-lactam-resistant *E. coli*. The results of the test are shown in the table below.

**Table 1.** MIC value of THES against several microorganisms.

| Microorganism |  | Microorganism |  |
| --- | --- | --- | --- |
| Gram-Positive Bacteria | MIC (%) | Gram-Negative Bacteria | MIC (%) |
| *B. subtilis* ATCC 6633 | 0.234 | *A. anitratus* | 0.115 |
| *S. aureus* ATCC 6538 | 0.234 | *E. coli* ATCC 25922 | 0.234 |
| *S. epidermidis* ATCC 12228 | 0.234 | *P. aeruginosa* ATCC 27853 | 0.115 |
| *C. diphtheria* | 0.810 | Carbapenem-resistant *K. pneumoniae* | 0.102 |
| *P. acne* | 0.200 | *S. typhi* | 0.115 |
| Vancomycin-resistant *E. faecalis* | 0.051 | β-lactam-resistant *E. coli* | 0.102 |
| Microorganism | MIC (%) |  |  |
| Fungi |  |  |  |
| *A. niger* | 0.937 |  |  |
| *C. albicans* ATCC 10231 | 0.469 |  |  |

### 4. TEM and SEM

*S. aureus*, a Gram-positive bacterium, and *E. coli,* a Gram-negative bacterium, were both observed using TEM (Fig. 5 and 6) and SEM (Fig. 7 and 8). It was shown that when THES was applied at 0.6%, some of the internal and external membranes were perforated, indicating extreme porosity of the internal and/or external cell membranes.

**Fig. 5.**
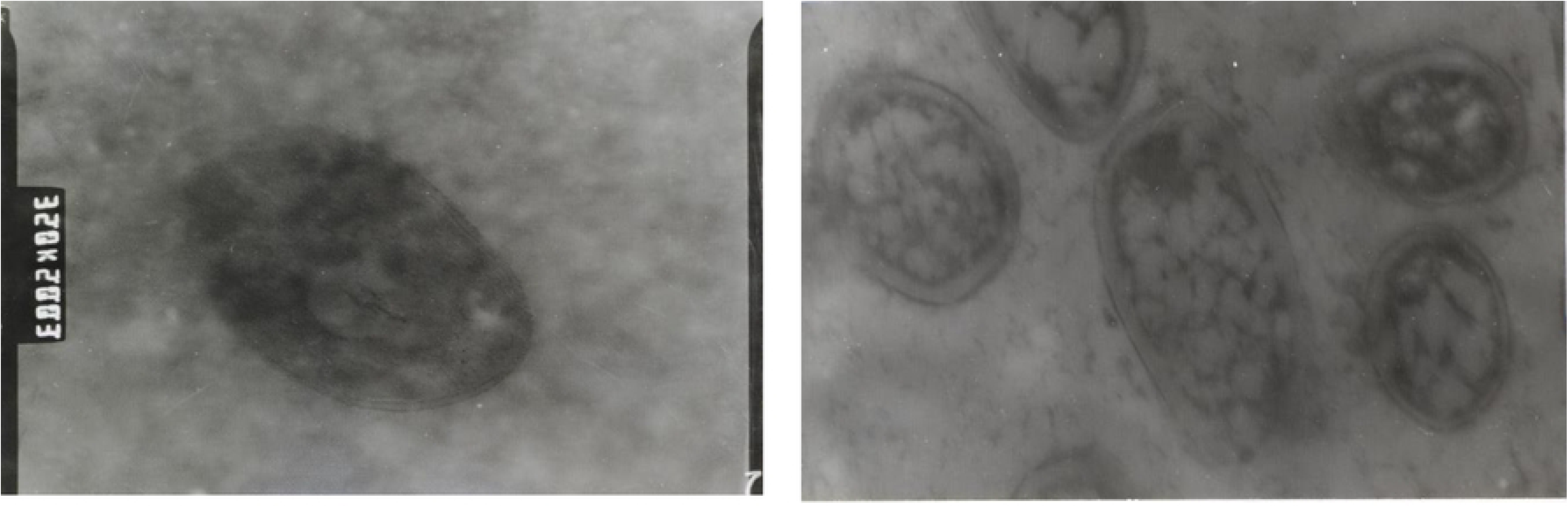
TEM result comparison of *S. aureus* under THES influence, normal cell and lysis cell (35,000× magnification).

**Fig. 6.**
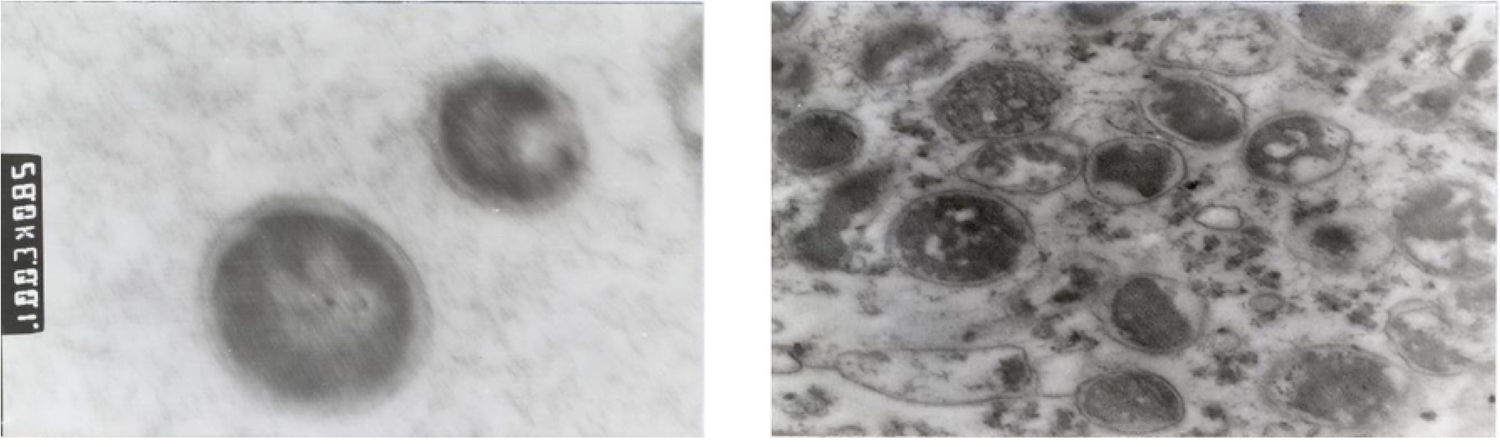
TEM result comparison of *E. coli* under THES influence, normal cell and lysis cell (35,000× magnification).

**Fig. 7.**
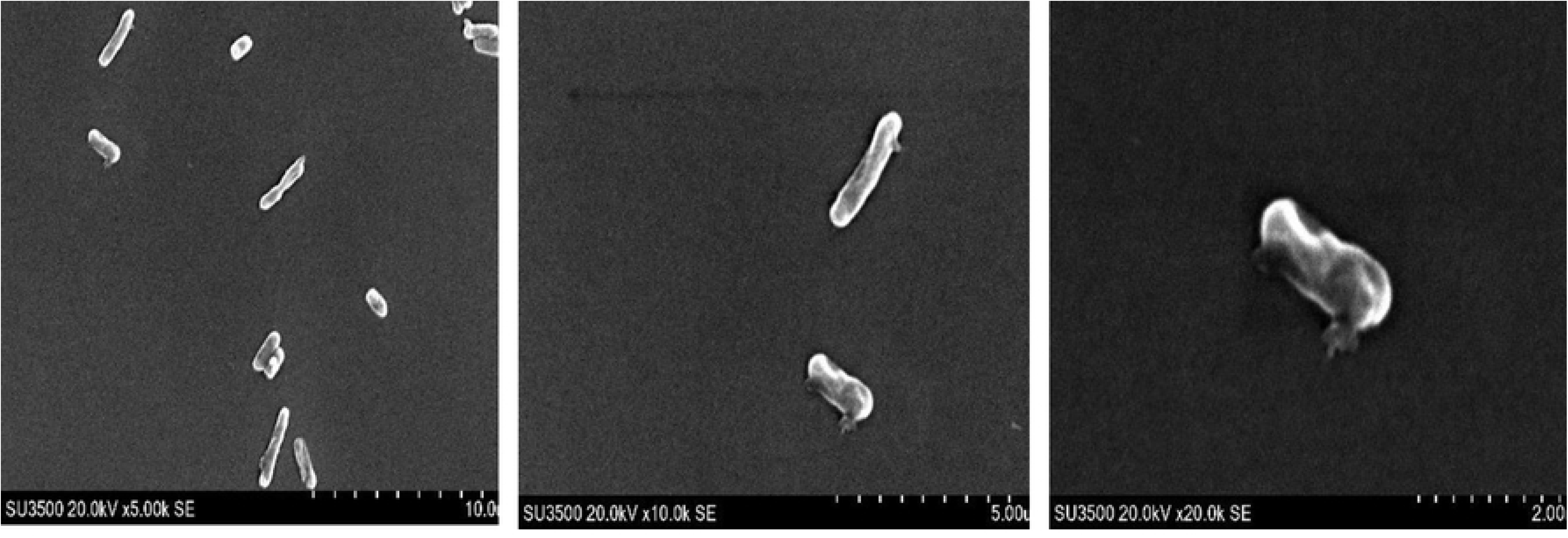
SEM observation of *S. aureus* under THES influence (5,000, 1,000, and 20,000× magnification).

**Fig. 8.**
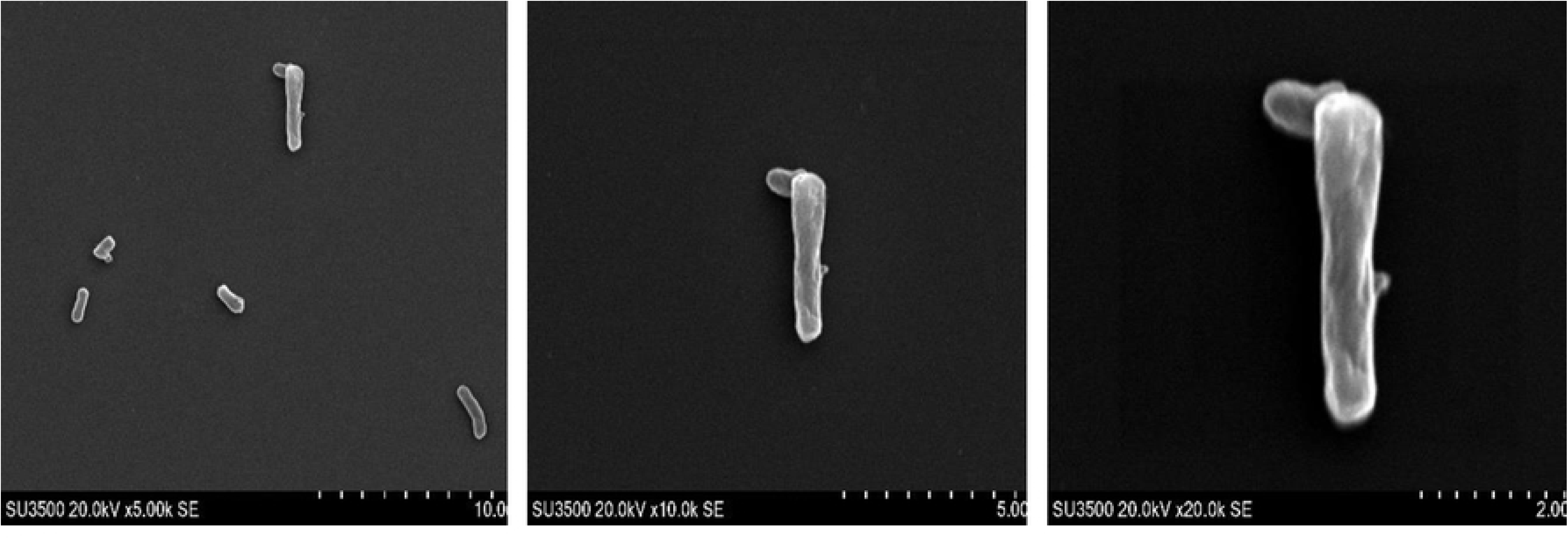
SEM observation of *E. coli* under THES influence (5,000, 10,000, and 20,000× magnification).

SEM observation results (Fig. 7 and 8) showed that both *S. aureus* and *E. coli* cell walls deformed significantly after being treated with THES, transforming from spherical to ellipsoidal, then into bar-like cells with a rough wrinkle membrane’s surface rather than a smooth membrane surface, and began leaking their cytoplasm.

### 5. AFM conferred the effect of THES morphology on the bacterial membrane

Fig. 9 and 10 showed the high-resolution images of the topography of bacteria specimens before and after THES treatment. Furthermore, Fig. 9b and 10b correspond to 3-dimensional images of bacteria. The images produced a visualization of a bacterial specimen with distinct features. The membranes of both bacterial specimens were deformed after treatment with THES. The differences in membrane surface structure were also noticed before and after treatment. The surface roughness of the bacterial wall changed significantly after treatment with THES.

**Fig. 9.**
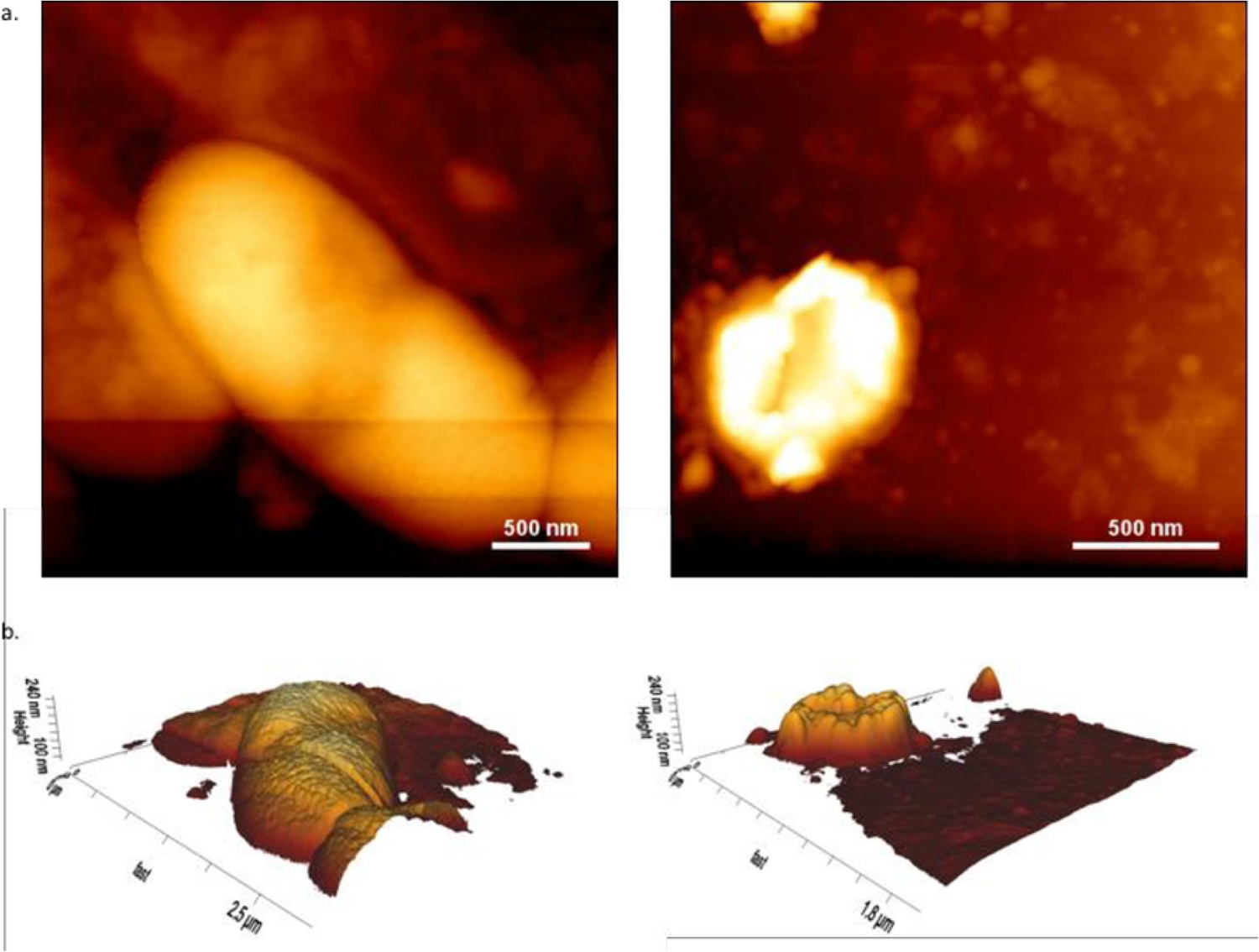
**AFM images of *E. coli***, where (a) corresponds to the topographic image of a 3 x 3 μm^2^ (left) and 2 x 2 μm^2^ (right) region of *E. coli* bacteria before and after, respectively; (b) represents the 3-dimensional topography of *E. coli* before (left) and after treatment (right) with the identical plane angle at 70°: 0°: 47° (X:Y:Z).

**Fig. 10.**
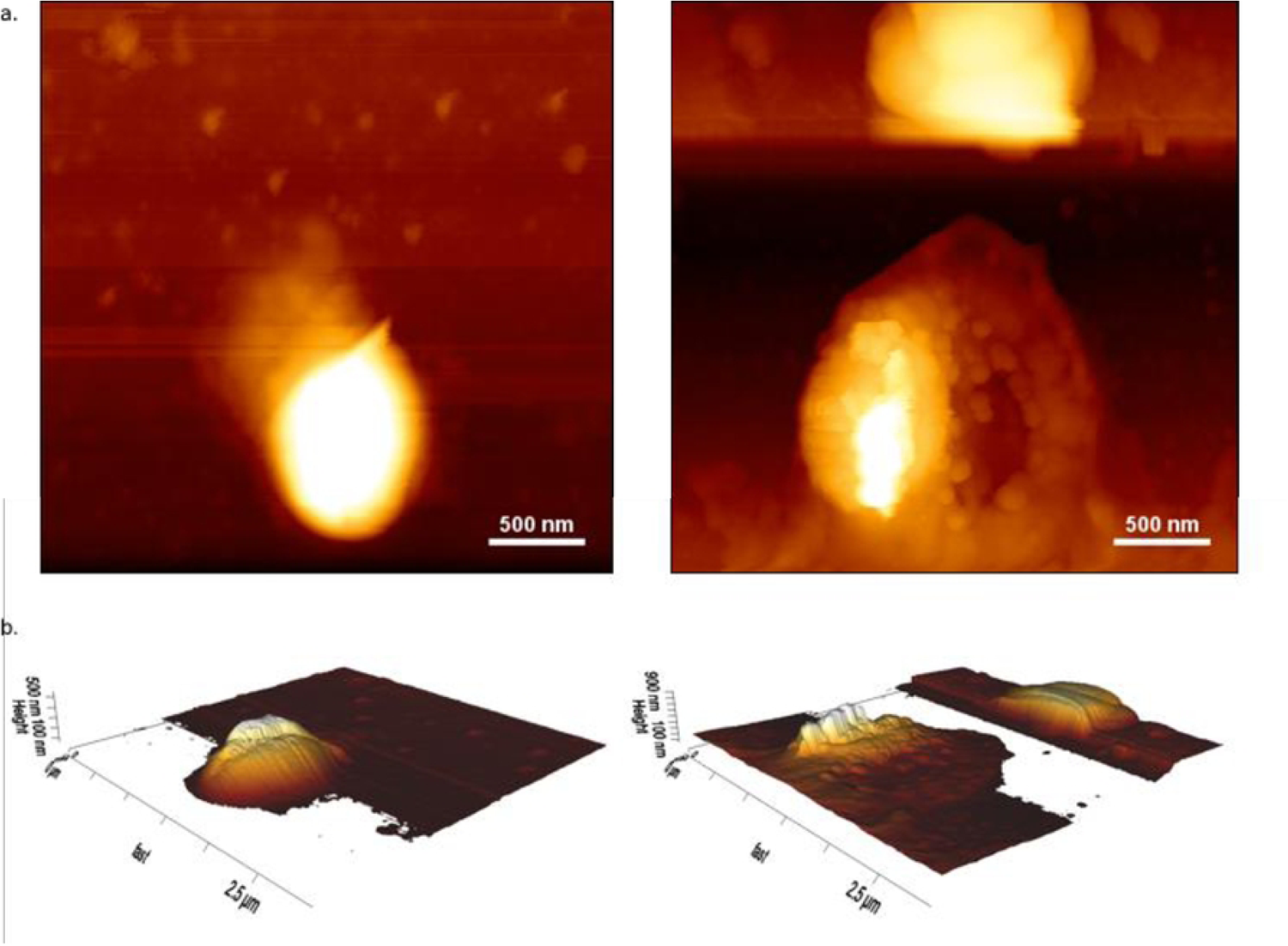
**AFM images of *S. aureus*,** where (a) corresponds to the topographic image of a 3 x 3 μm^2^ (left) and 3 x 3 μm^2^ (right) region of *S. aureus* bacteria before and after, respectively; (b) represents 3-dimensional topography of *S. aureus* before (left) and after treatment (right) with the identical plane angle at 70°: 0°: 47° (X:Y:Z).

Fig. 11 and 12 correspond to wide and length cross-section measurements of *E. coli* and *S. aureus* respectively, before and after THES treatment. Before treatment, the histogram plot measurement of *E. coli* in Fig. 11 had a maximum height of approximately 80 nm, with a smooth curve projection with gradual rising and falling, demonstrating the integrity of cell plasma contained by a membrane. In contrast, the maximum height of treated *E. coli* was observed to be around 110 nm. This was followed by a valley-like structure in the topography of the cell membrane. Height fluctuations were observed starting from −20 nm, rising to 110 nm, followed by a descending to 60 nm, a slight increase to 80 nm, and went down to 0. This pattern was further observed in the cross-section plot (Fig. 11). This is in contrast to the histogram plot of *E. coli* after treatment with THES. The treated specimen displayed a valley-like structure, fluctuating from 5 nm, reaching its peak at 100 nm, sinking to 50 nm, followed by a rise to 80 nm, and descending to 0. Histogram plot recorded change in the maximum height of *E. coli* species before and after treatment with THES. The increase in height might be induced by the deformation of the bacterial cell wall. However, factors that limit and induce bacterial cell wall deformation were not measured in the study. It is assumed that components and the valley-like structure are the projection of released cell plasma and organelles, due to the disruption of the cell wall by THES.

**Fig. 11.**
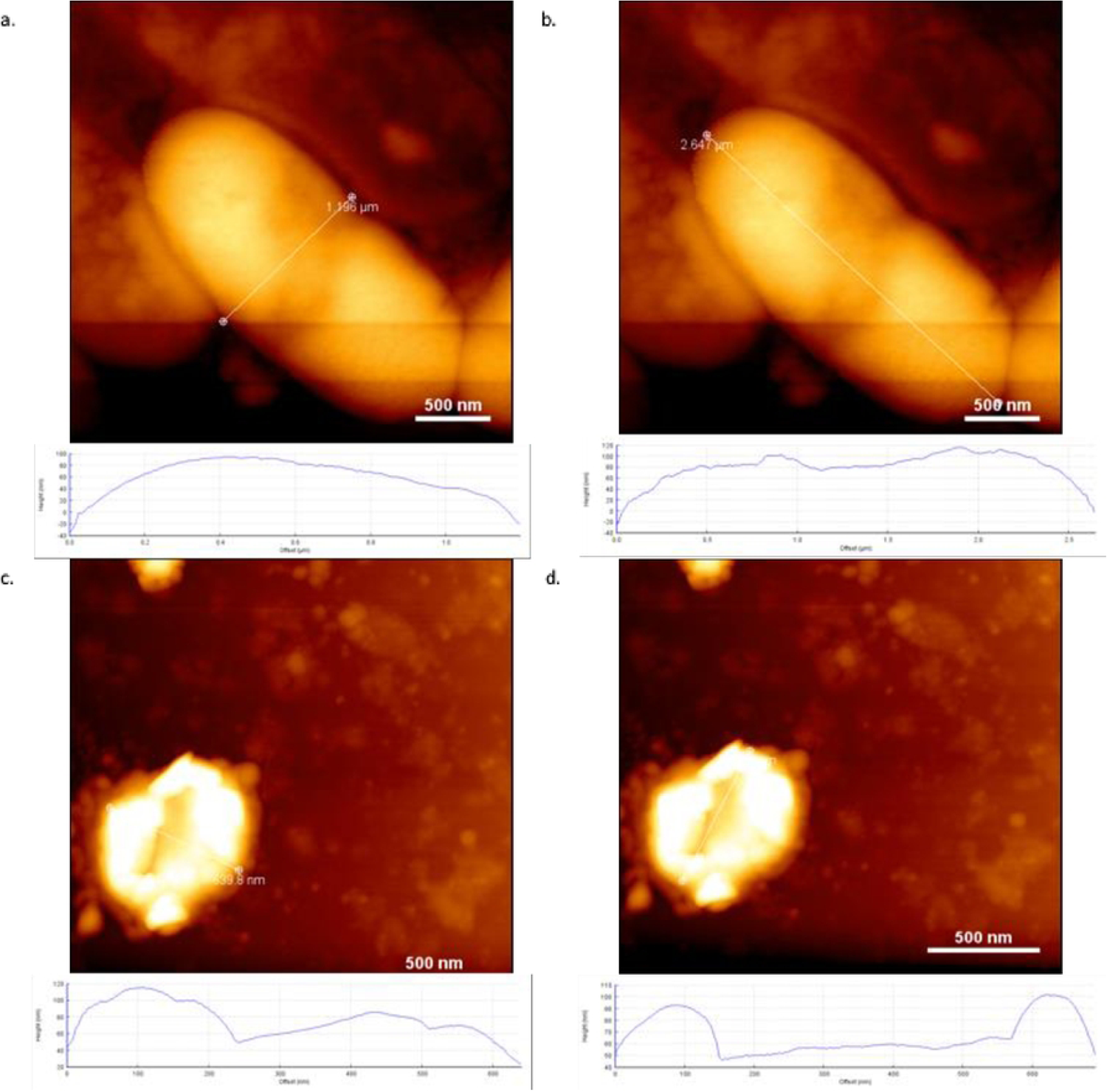
**A plot of cross-section measurement of *E. coli*** in width cross-section (a, c) and length cross-section (b, d). Upper images show *E. coli* before treatment, whereas lower images were taken after treatment with the THES.

**Fig. 12.**
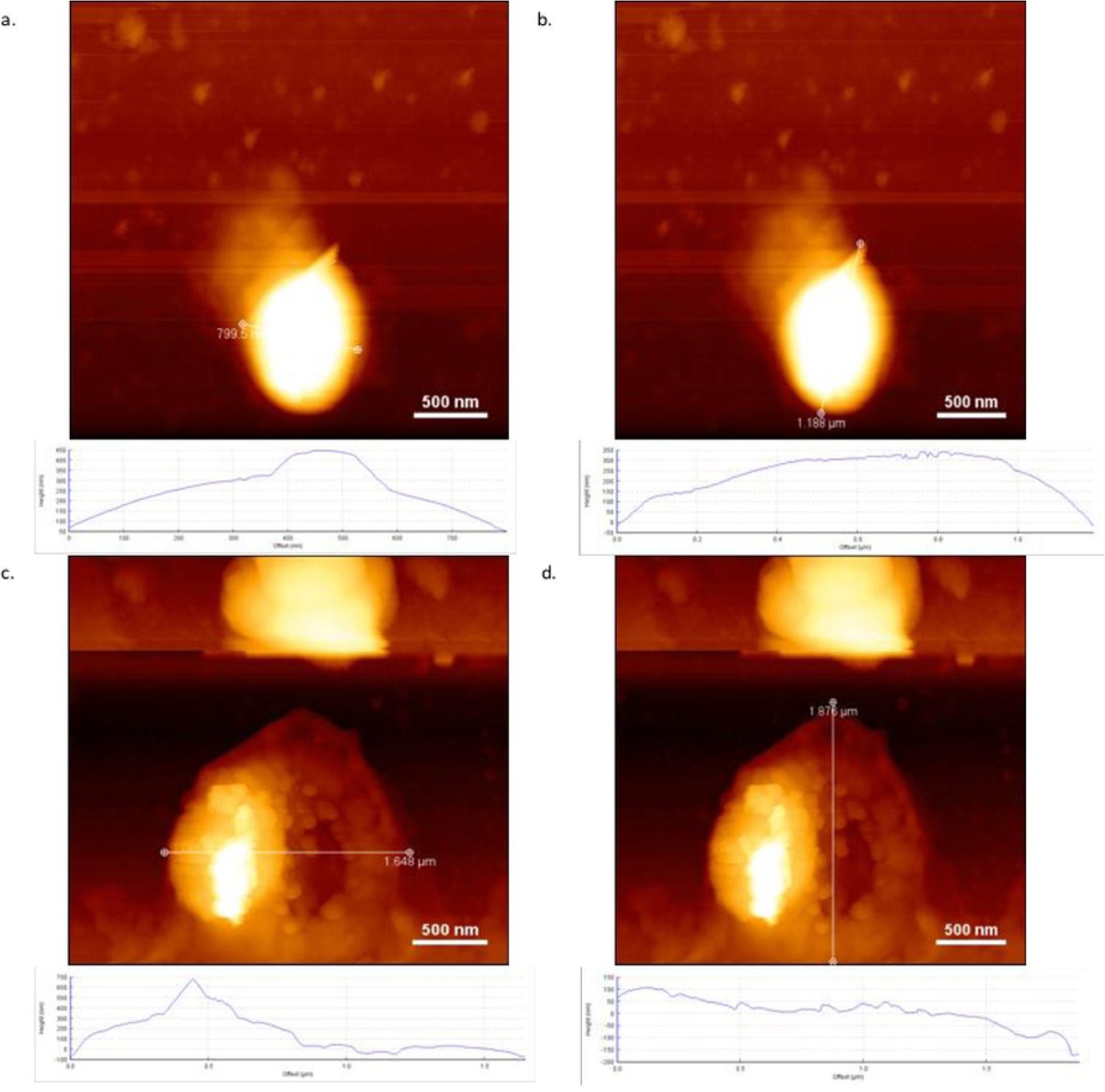
**A plot of cross-section measurement of *S. aureus*** in width cross-section (a, c) and length cross-section (b, d). Upper images show *S. aureus* before treatment, whereas lower images were taken after treatment with THES.

Fig. 12a and 12c correspond to the wide cross-section measurement of *S. aureus* specimen, whereas Fig. 12b and 12d correspond to the length cross-section measurement before (upper images) and after THES treatment (upper images). A similar pattern of deformation with *E. coli* was observed in both planes. A smooth curvature structure was exhibited by *S. aureus* before treatment with THES. The structure was then replaced with a sharp valley-like structure after treatment with the THES. However, in *S. aureus*, fluctuation of the valley-like structures is more defined and with a higher frequency of spikes observed. The most observable difference observed is the significant decrease in height in both the width and length plane of *S. aureus* after being treated with THES. Before treatment, the maximum height of the specimen was observed at 450 nm and the minimum height was at 50 nm. In the treated specimen, the maximum height was recorded at 500 nm and the minimum height was at −100 nm.

Significant difference observed in the plane of *S. aureus* before and after treatment (Fig. 9 and 12) was assumed to be caused by an uneven distribution of poly-L-lysine during the glass slide coating step. The increase in plane height induced by the utilization of poly-L-lysine as an immobilization agent was studied [40] and recorded at approximately 40–50 nm. The break-line artifact observed during AFM imaging could be caused by the nature of the sample and the mode used during the experiment. Furthermore, high-resolution AFM images of the membrane may result in overestimation due to the radius of curvature of the AFM tip [41].

### 6. Peptidoglycan Binding Test of THES

This test was performed to show that THES, as an antibacterial agent postulated to work in cell membranes composed of protein & peptidoglycan, was indeed affected by either protein or peptidoglycan. The test was conducted at 25°C with 30 µM of peptidoglycan sample and 380 µM of THES using MicroCal ITC200 (MicroCal Inc., North Hampton, MA). The ITC sigmoidal curve results indicated that THES formed multiple binding sites with peptidoglycan with ΔH = −25.730 ± 3.246 kcal/mol (Fig. 13). The calculated dissociation constant value (*K*_*d*_) was 17.5 ± 73.5 nM. This *K*_*d*_ was considered a relatively tight interaction [38]. This supported the hypothesis that THES binds to peptidoglycan as a chelating agent. The curve results for negative control using BSA (*K*_*d*_ = 2.62 nM) showed no indication of binding between BSA and THES, suggesting that THES would not non-specifically bind with most proteins (Fig. 14). The results were also consistent with the *in silico* molecular docking between THES and peptidoglycan conducted in this study in which THES binds to four different residues of peptidoglycan (Fig. 4).

**Fig. 13.**
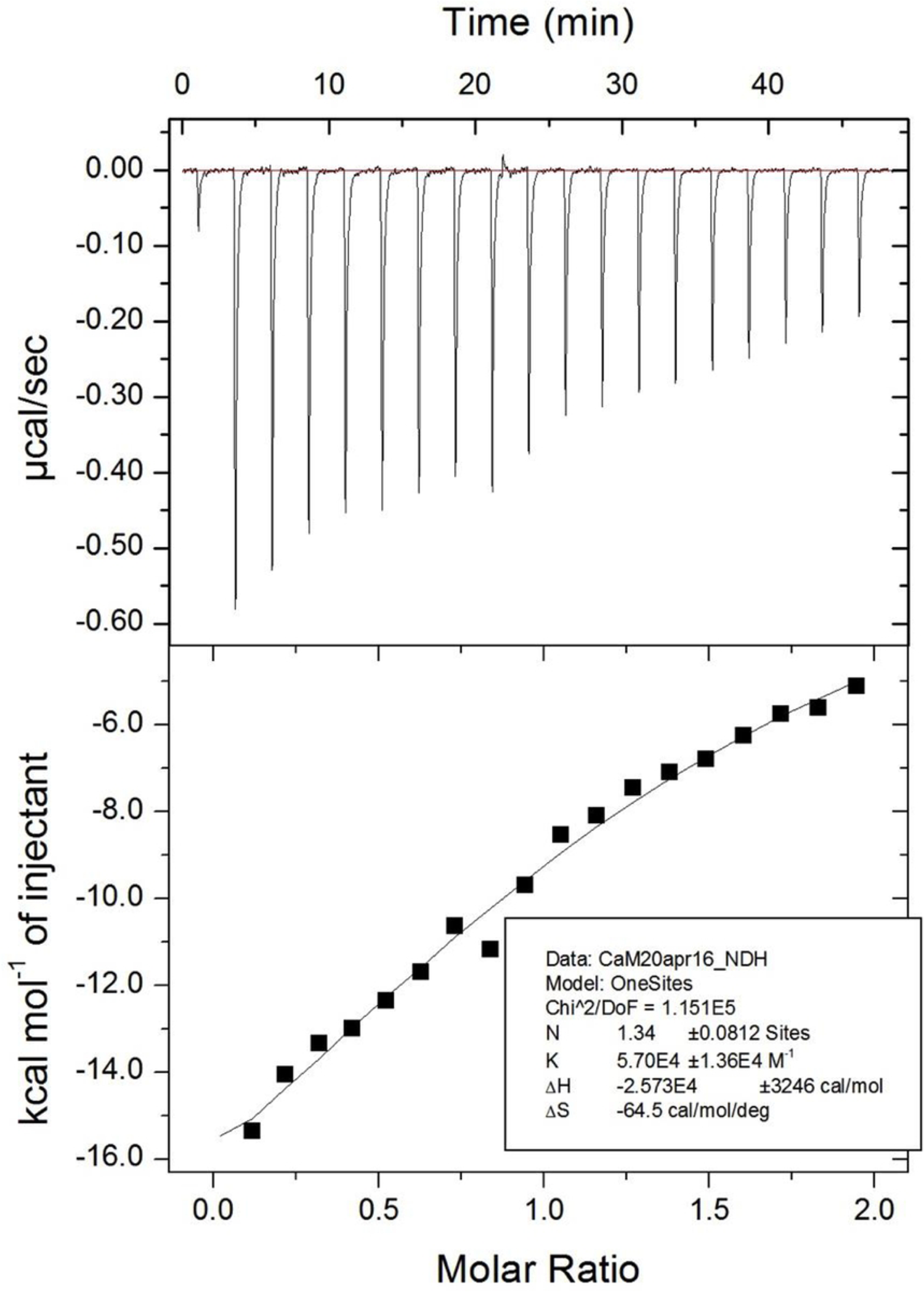
Result of glycan binding test of THES.

**Fig. 14.**
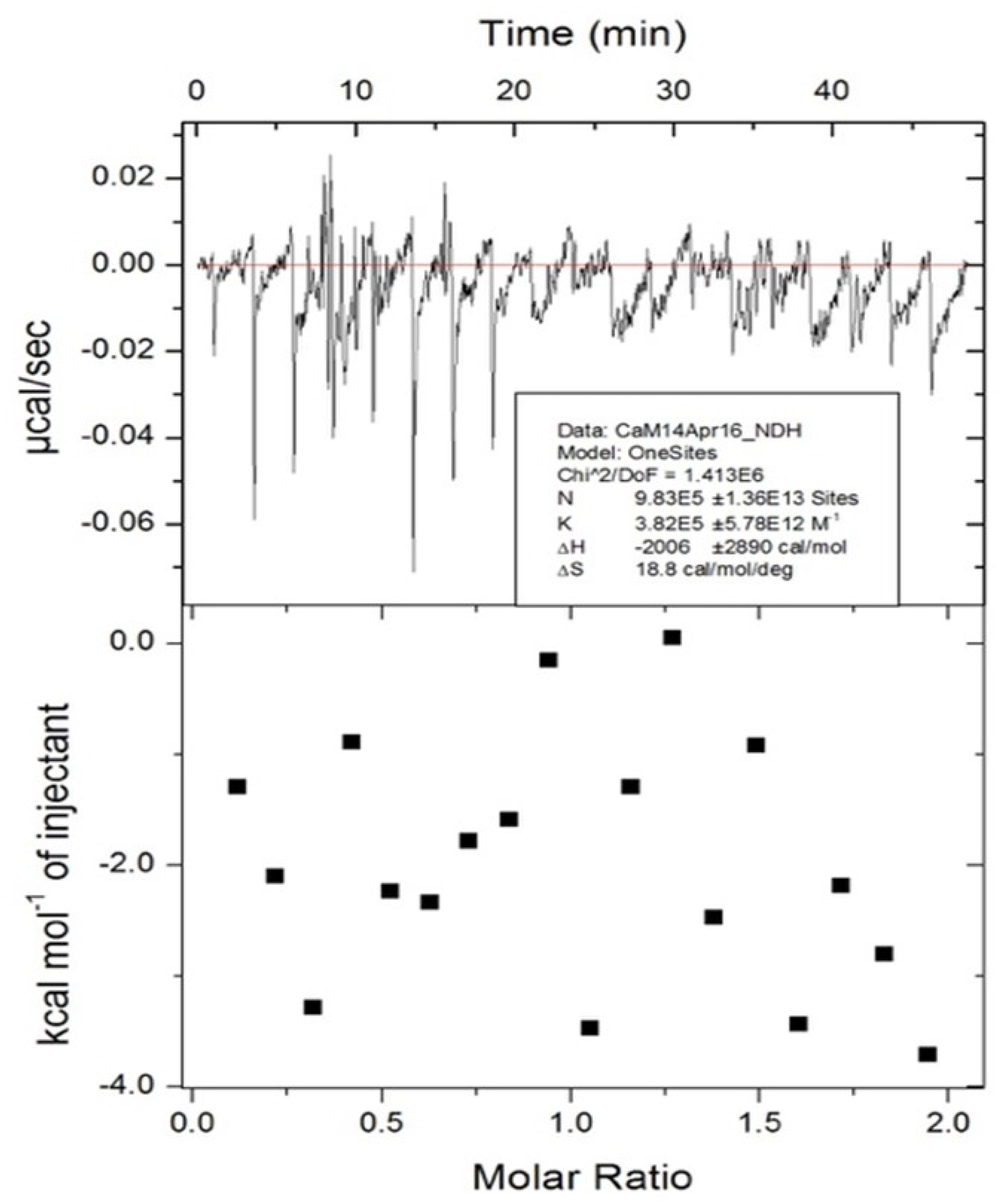
Result of glycan binding test of BSA.

### 7. Resistance Test

The MIC and phenol coefficient of THES against *S. aureus* were plotted in an integrated bar-line graph, respectively, versus the period of THES treatment (Fig. 15). MIC dosage at contact time 10 minutes for phenol was 7.69 mg/mL, which was at a dilution ratio of 1:130; whereas THES treatment throughout the period resulted in overall MIC at 7.69, 8.01, and 8.33 mg/mL, which was at dilution ratio 1:30, 1:125, and 1:120 respectively. The phenol coefficients of THES treated at all seven months against *S. aureus* were relatively stable at approximately 92-100%, showing considerable antiseptic activity even comparable to the positive control phenol. Over seven months, the phenol coefficient of THES did not decline, indicating that the compound was not prone to *S. aureus* resistance.

**Fig. 15.**
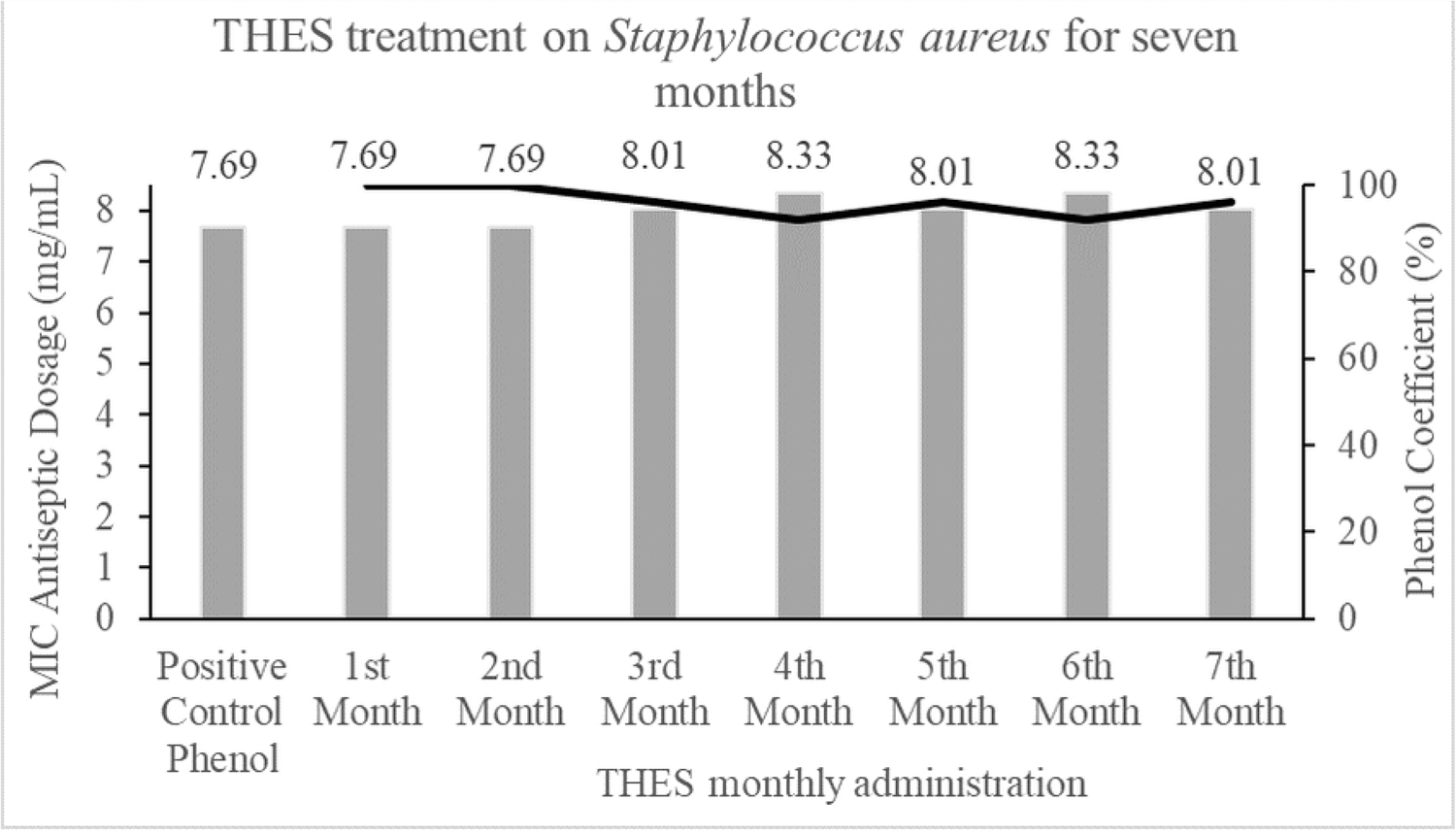
Trend on MIC dosage and phenol coefficient of THES for seven months treatment period on *S. aureus*.

## Discussions

An effective antibacterial agent has two important characteristics. First, it should be either bacteriostatic or bactericidal. Second, they are typically molecules and/or compounds that can be manipulated and modified to identify analogues, with desirable antibacterial activities against the desired range of target pathogens and to achieve good PK/PD properties with acceptable low levels of toxicity against mammalian cells [42–44]. In addition to these, the most valuable antibiotics are soluble drugs that are chemically stable if care is taken orally with systemic effect [20,42].

THES is a chelating agent, characterized by its strong sulfate active group, providing it with the capability to bind strongly into metal, inorganic or organic groups/ligands. Greater chelation capability or higher electronegativity will result in stronger interference to the cross-linking bond of protein or peptidoglycan and similar types of polymer. Additionally, the more electronegative the X group (Fig.16), the stronger the chelating agent and/or the stronger the bond; such as sulfate is stronger than phosphate, hydroxy, carboxylate, and the other organic groups. On the contrary, the more electronegative the X group, the lower the pH solution. In the case of more inorganic groups, pressure and temperature will strengthen the internal bound structure. This meant that the polymer with the more inorganic group will withstand higher temperatures and pressure, which means it is more suitable for high-temperature and high-pressure applications.

**Fig. 16.**
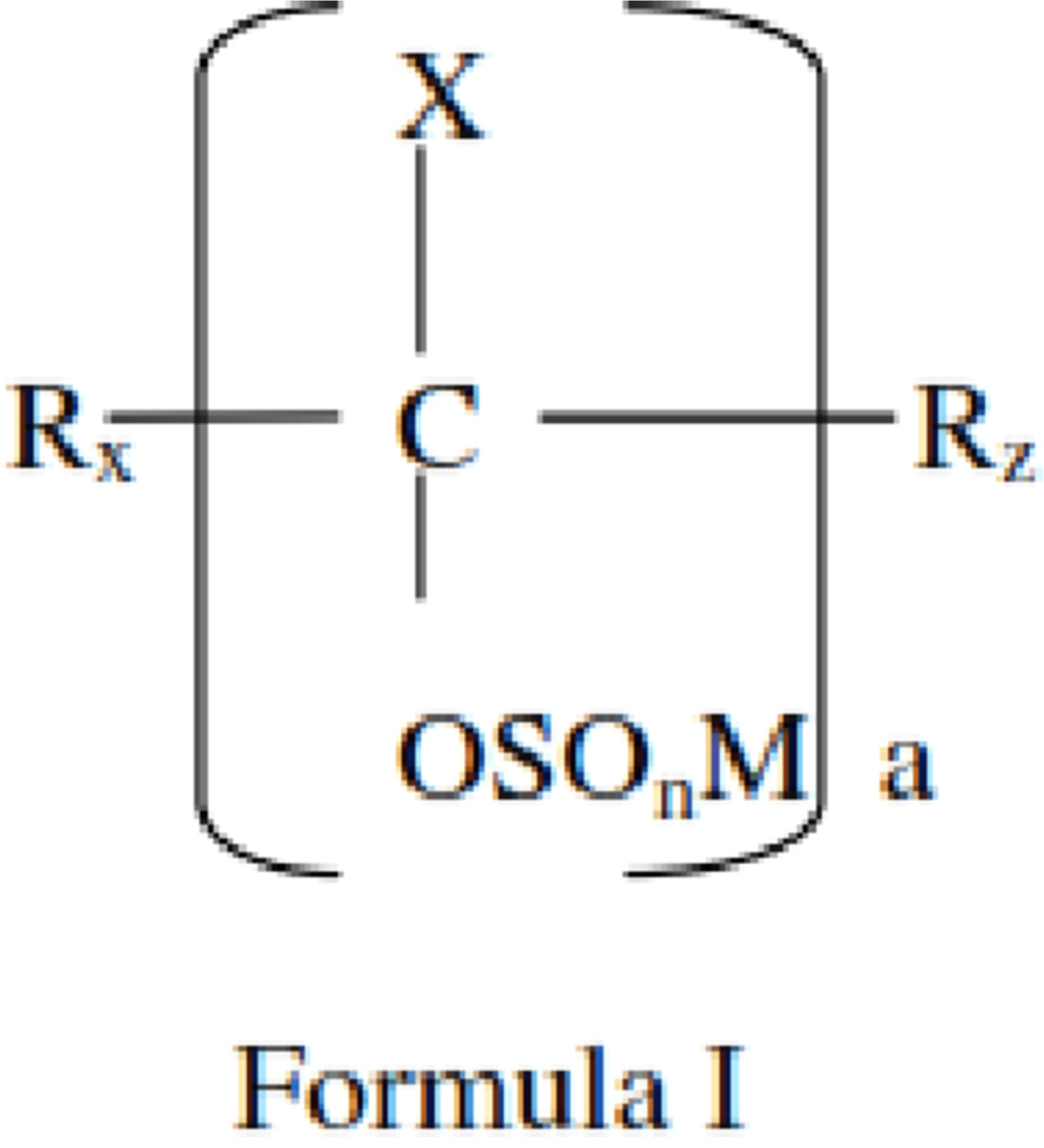
Structure of THES.

To a further extent, a sudden change in sample height might affect the geometric model captured by the AFM instrument. Experiments conducted should have monitored the pre-conditioning of bacteria as a sample. This includes possibilities of contaminating substrata on slides upon application of immobilization agent, such as debris and dust, as well as exchange and calibration of cantilever tip. Differences in characteristics projected by specimens *E. coli* and *S. aureus* might be attributed to the different structures of the bacterial wall of the bacteria. *E. coli* as a Gram-negative bacterium is characterized by its single layer of peptidoglycan. In contrast, *S. aureus* as a Gram-positive bacterium is characterized by multiple layers of peptidoglycan. The result data illustrates that the overall size of *E. coli* specimens after treatment with THES showed a significant decrease, compared with *E. coli* under normal conditions [45]. On the other hand, *S. aureus* specimens treated with THES event seemed to undergo expansion due to a burst-like event. The treated specimen was observed to occupy an area that was larger by 1.5x than its normal size [46]. Hence, *S. aureus* was assumed to show a more reactive interaction when treated with THES.

The data obtained in the study proved that the THES mechanism resulted in a disruption event on the bacterial cell membrane. The visual evidence observed supported the proposed mechanism of THES. In comparison to live-dead staining using both conventional and confocal laser scanning microscopes, this method has demonstrated a high potential for direct observation of live cells in air mode (without liquid medium) for drug testing and development. The study benefited from AFM visualization in several planes, which provided sufficient evidence of bacterial membrane disruption as a mechanism of THES as a non-resistant antibacterial agent.

As a new antibacterial, THES showed all the features mentioned above. THES is a bactericidal agent that is effective against Gram-positives such as *S. aureus*, Gram-negative such as *E. coli*, and fungi such as *Aspergillus niger*. Bacterial action involves interaction with the peptidoglycan cell membrane. THES targets the cross-linking bond among peptides in the protein structure or glycan in the peptidoglycan structure, loosening it, and resulting in an increase in the porosity of the peptidoglycan cell membrane and cell lysis, which leads to cell death. This interference of THES into the cross-linking bond of protein or peptidoglycan can range from slightly loosening to completely cutting the cross-linking bond, depending on the concentration of THES in the system.

THES had a solubility of 31.5% at room temperature and the higher temperature will increase the solubility. Solubility and permeability assessments are crucial for ruling in or out the potential of a compound to be a drug. Drug substance often needs access to a patient’s circulation and therefore may be injected or more generally have to be absorbed in the digestive system [47,48]. *In vitro* activity study of THES against several antibiotic-resistant bacteria such as vancomycin-resistant *E. faecalis*, β-Lactam resistant *E. coli,* and carbapenem-resistant *K. pneumonia* showed considerably very low MIC values between 0.05-0.1%.

Further following a seven-month resistance study, THES-treated *S. aureus* also exhibited little-to-no MDR characteristics. In addition, THES exposure also resulted in stable antibacterial characteristics comparable to phenol, indicating that this novel compound is also potentially effective as a disinfectant against specific bacteria. As demonstrated earlier via *in silico* and *in vitro* characterization, the suggested chelation between the compound and peptidoglycan layers made THES an effective long-term antiseptic and drug. The test showed that even after being grown at a 0.3% MIC dosage of THES for seven months, *S. aureus*, a Gram-positive bacteria, could not adapt and counteract the underlying peptidoglycan-targeting properties. This also made it a highly sought-after clinical antiseptic as *S. aureus* can easily acquire MDR characteristics, allowing it to turn into variants such as MRSA [49,50].

## Conclusions

Evaluation of the tetra hydroxy ethyl disulfate disodium (THES) as a potential non-resistant antibacterial chelating agent had been conducted through *in silico* and *in vitro* experiments. THES had been tested and proven effective against several bacterial strains with a relatively low (0.05-0.1%) minimum inhibitory concentration. THES was also found to express antibacterial properties after treatment on *Staphylococcus aureus* during the seven-month resistance study.

Bacterial cell morphology analyses using different microscopy instrumentations strongly indicated that the administration of THES causes bacterial cell lysis, leading to cell death. The hypothesis was that THES bound to the cell membrane with peptidoglycan as its target. An *in silico* study confirmed that THES binds to four different residues of the peptidoglycan structure. This result was further supported by an *in vitro* binding study between THES with peptidoglycan molecules using the isothermal titration calorimetry. Taken altogether, THES has shown characteristics as an effective antibacterial agent with a mode of action as a chelating agent that binds to peptidoglycan with little observable effect on the host. As a result of this binding mechanism, THES has potential as a novel non-resistant antibacterial chelating agent.

## Acknowledgments

The authors would like to express our highest gratitude to PT. Novis Natura Navita, the late Mr. Haryanto Wardoyo, and Mr. Bobby Hadipraja who had provided research grants awarded to Kholis Abdurachim Audah with Grant Number: AG/SGU/Coop/0018/VI/2016. The authors also thank the Swiss German University for financial support through the Central Research Fund and the Faculty Research Fund to Kholis Abdurachim Audah. This article is submitted in memory of the departure of one of the funders, the late Mr. Haryanto Wardoyo, our coauthor the late Dr. Doddy Kustaryono (STKIP Surya), and the late Dr. Dedy HB Wicaksono (Swiss German University) who helped us bridge the collaboration between the Swiss German University and the University of Technology Malaysia. Thanks to all who have helped with the research activities either directly or indirectly.

## Supporting Information

**No supporting information was appended.**

## References

1. Norris V, Molina F, Gewirtzc AT. Hypothesis: Bacteria control host appetites. J Bacteriol. 2013;195: 411–416. doi:10.1128/JB.01384-12

2. Laxminarayan R, A D, C W, A Z, H W. Antibiotic resistance-the need for global solutions. Lancet Infect Dis. 2013;13: 1057–98.

3. Ventola CL. The Antibiotic Resistance Crisis. P&T. 2015;40: 278–283. doi:10.5796/electrochemistry.82.749

4. Crnich CJ, Jump R, Trautner B, Sloane PD, Mody L. Optimizing Antibiotic Stewardship in Nursing Homes: A Narrative Review and Recommendations for Improvement. Drugs and Aging. Springer International Publishing. 2015;32: 699–716. doi:10.1007/s40266-015-0292-7

5. Sophie T, Drissner D, Walsh F. Antimicrobial Resistance in Agriculture. Am Soc Microbiol. 2016;7: e02227–15. doi:10.1128/mBio.02227-15.

6. Huang SS, Chen CL, Huang FW, Johnson FE, Huang JS. Ethanol Enhances TGF-β Activity by Recruiting TGF-β Receptors from Intracellular Vesicles/Lipid Rafts/Caveolae to Non-Lipid Raft Microdomains. J Cell Biochem. 2016;117: 860–871. doi:10.1002/jcb.25389

7. Kathleen MM, Samuel L, Felecia C, Reagan EL, Kasing A, Lesley M, et al. Antibiotic Resistance of Diverse Bacteria from Aquaculture in Borneo. Int J Microbiol. 2016;2016. doi:10.1155/2016/2164761

8. Haag LM, Fischer A, Otto B, Plickert R, Kühl AA, Göbel UB, et al. Intestinal microbiota shifts towards elevated commensal escherichia coli loads abrogate colonization resistance against campylobacter jejuni in mice. PLoS One. 2012;7: 1–13. doi:10.1371/journal.pone.0035988

9. Mobegi FM, van Hijum SAFT, Burghout P, Bootsma HJ, de Vries SPW, van der Gaast-de Jongh CE, et al. From microbial gene essentiality to novel antimicrobial drug targets. BMC Genomics. 2014;15. doi:10.1186/1471-2164-15-958

10. Mandal SM, Roy A, Ghosh AK, Hazra TK, Basak A, Franco OL. Challenges and future prospects of antibiotic therapy: From peptides to phages utilization. Front Pharmacol. 2014 may 5. pii: 1–12. doi:10.3389/fphar.2014.00105.

10. Reygaert WC. An overview of the antimicrobial resistance mechanisms of bacteria. AIMS Microbiology. 2018;4(3):482–501. doi: 10.3934/microbiol.2018.3.482

11. Baym M, Stone LK, Kishony R. Multidrug evolutionary strategies to reverse antibiotic resistance. Science. 2015;351(6268):3292– 3292. doi:10.1126/science.aad3292

12. Barlam TF, Cosgrove SE, Abbo LM, Macdougall C, Schuetz AN, Septimus EJ, et al. Executive summary: Implementing an antibiotic stewardship program: Guidelines by the infectious diseases society of America and the society for healthcare epidemiology of America. Clin Infect Dis. 2016;62: 1197–1202. doi:10.1093/cid/ciw217

13. Cieplik F, Deng D, Crielaard W, Buchalla W, Hellwig E, Al-Ahmad A, Maisch T. Antimicrobial photodynamic therapy – what we know and what we don’t. Critical Reviews in Microbiology. 2018;44(5):571–589. doi:10.1080/1040841x.2018.1467876

14. Rutala WA, Weber DJ. Disinfection and sterilization: An overview. Am J Infect Control. Elsevier Inc. 2013;41: S2–S5. doi:10.1016/j.ajic.2012.11.005.

15. Barah P, Winge P, Kusnierczyk A, Tran DH, Bones AM. Molecular Signatures in Arabidopsis thaliana in Response to Insect Attack and Bacterial Infection. PLoS One. 2013;8. doi:10.1371/journal.pone.0058987

16. Costin BN, Miles MF. Molecular and neurologic responses to chronic alcohol use. Alcohol and the Nervous System. 2014; 157–171. doi:10.1016/b978-0-444-62619-6.00010-0

17. Tamar Ringel-Kulka CHC, Temas D, Kim A, Maier DM, Scott K, Galanko JA, et al. Altered Colonic Bacterial Fermentation as a Potential Pathophysiological Factor in Irritable Bowel Syndrome. Am J Gastroenterol. 2015;110: 1339–1346.

18. León-Buitimea A, Garza-Cárdenas CR, Garza-Cervantes JA, Lerma-Escalera JA, Morones-Ramírez JR. The Demand for New Antibiotics: Antimicrobial Peptides, Nanoparticles, and Combinatorial Therapies as Future Strategies in Antibacterial Agent Design. Frontiers in Microbiology. 2020;11:1–10. doi:10.3389/fmicb.2020.01669

19. Hughes D, Karlén A. Discovery and preclinical development of new antibiotics. Ups J Med Sci. 2014;119: 162–169. doi:10.3109/03009734.2014.896437

20. Balouiri M, Sadiki M, Ibnsouda SK. Methods for in vitro evaluating antimicrobial activity: A review. J Pharm Anal. Elsevier; 2016;6: 71–79.

21. Velkov T, Roberts KD, Nation RL, Thompson PE, Li J. Pharmacology of polymyxins: new insights into an ‘old’ class of antibiotics. Int J Ambient Comput Intell. 2013;8. doi:10.2217/fmb.13.39.Pharmacology

22. Schechner V, Temkin E, Harbarth S, Carmeli Y, Schwaber MJ. Epidemiological interpretation of studies examining the effect of antibiotic usage on resistance. Clin Microbiol Rev. 2013;26: 289–307. doi:10.1128/CMR.00001-13

23. Tacconelli E, Cataldo MA, Paul M, Leibovici L, Kluytmans J, Schröder W, et al. STROBE-AMS: Recommendations to optimise reporting of epidemiological studies on antimicrobial resistance and informing improvement in antimicrobial stewardship. BMJ Open. 2016;6: 1–8. doi:10.1136/bmjopen-2015-010134

24. Verhoef TI, Morris S. Cost-effectiveness and Pricing of Antibacterial Drugs. Chemical Biology & Drug Design. 2014;85(1):4–13. doi:10.1111/cbdd.12417

25. Pirhadi S, Sunseri J, Koes DR. Open source molecular modeling. Journal of Molecular Graphics and Modelling. 2016;69:127–143. doi:10.1016/j.jmgm.2016.07.008

26. Li H, Leung KS, Wong MH, Ballester PJ. Improving AutoDock Vina Using Random Forest: The Growing Accuracy of Binding Affinity Prediction by the Effective Exploitation of Larger Data Sets. Molecular. Informatics. 2015;34(2-3):115–126. doi:10.1002/minf.201400132.

27. Gaillard T. Evaluation of AutoDock and AutoDock Vina on the CASF-2013 Benchmark. Journal of Chemical Information and Modeling. 2018;1–30 doi:10.1021/acs.jcim.8b00312

28. Stylianou A. Atomic Force Microscopy for Collagen-Based Nanobiomaterials. Journal of Nanomaterials. 2017;1–14. doi:10.1155/2017/9234627

29. Dufrêne YF, Ando T, Garcia R, Alsteens D, Martinez-martin D, Engel A, et al. Imaging modes of atomic force microscopy for application in molecular and cell biology. Nat Publ Gr. Nature Publishing Group; 2017;12: 295–307. doi:10.1038/nnano.2017.45

30. Vahabi S. Atomic Force Microscopy Application in Biological Research: A Review Study. 2013;38.

31. Müller DJ, Dumitru AC, Lo Giudice C, Gaub HE, Hinterdorfer P, Hummer G, Alsteens D. Atomic Force Microscopy-Based Force Spectroscopy and Multiparametric Imaging of Biomolecular and Cellular Systems. Chemical Reviews. 2020. doi:10.1021/acs.chemrev.0c00617

32. Huang Q, Wu H, Cai P, Fein JB, Chen W. Atomic force microscopy measurements of bacterial adhesion and biofilm formation onto clay-sized particles. Nat Publ Gr. Nature Publishing Group. 2015; 1–12. doi:10.1038/srep16857.

33. Wu S, Zuber F, Maniura-Weber K, Brugger J, Ren Q. Nanostructured surface topographies have an effect on bactericidal activity. Journal of Nanobiotechnology. 2018;16(1). doi:10.1186/s12951-018-0347-0

34. Mukherjee R, Saha M, Routray A, Chakraborty C. Nanoscale Surface Characterization of Human Erythrocytes by Atomic Force Microscopy: A Critical Review. IEEE Transactions on NanoBioscience. 2015;14(6):625–633. doi:10.1109/tnb.2015.2424674

35. Hyldgaar M, Mygind T, Vad BS, Stenvang M, Otzen DE, Meyer RL. The Antimicrobial Mechanism of Action of Epsilon-Poly-l-Lysine. Applied and Environmental Microbiology. 2014;80(24):7758–7770. doi:10.1128/aem.02204-14

36. Sidiq S, Prasad GVRK, Mukhopadhaya A, Pal SK. Poly(l-lysine)-Coated Liquid Crystal Droplets for Cell-Based Sensing Applications. The Journal of Physical Chemistry B. 2017;121(16):4247–4256. doi:10.1021/acs.jpcb.7b00551

37. Audah K. Fundamentals of protein-protein interactions and their methods of characterization. Bioinformatics and Biomedical Research Journal. 2021;4(2):70–82. doi: 10.11594/bbrj.04.02.04

38. England JW. The phenol coefficient method of testing disinfectants. Journal of the American Pharmacists Association. 2006;2(8):955–958.

39. Li M, Wu H, Wang Y, Yin T, Gregersen H, Zhang X, Wang G. Immobilization of heparin/poly-l -lysine microspheres on medical grade high nitrogen nickel-free austenitic stainless steel surface to improve the biocompatibility and suppress thrombosis. Materials Science and Engineering: C. 2017;73:198–205. doi:10.1016/j.msec.2016.12.070

40. Vorselen D, Piontek MC, Roos WH, Wuite GJL. Mechanical Characterization of Liposomes and Extracellular Vesicles, a Protocol. Frontiers in Molecular Biosciences. 2020;7;1–14. doi:10.3389/fmolb.2020.00139

41. Nemeth J, Oesch G, Kuster SP. Bacteriostatic versus bactericidal antibiotics for patients with serious bacterial infections: systematic review and meta-analysis. Journal of Antimicrobial Chemotherapy. 2015;70(2):382–395. doi:10.1093/jac/dku379

42. Bull SC, Doig AJ. Properties of protein drug target classes. PLoS One. 2015;10: 1–44. doi:10.1371/journal.pone.0117955

43. Vatsraj S, Chauhan K, Pathak H. Formulation of a novel nanoemulsion system for enhanced solubility of a sparingly water soluble antibiotic, clarithromycin. J Nanosci. 2014;2014: 1–7. doi:10.1155/2014/268293

44. Gumbart JC, Beeby M, Jensen GJ. Escherichia coli Peptidoglycan Structure and Mechanics as Predicted by Atomic-Scale Simulations. 2014;10. doi:10.1371/journal.pcbi.1003475.

45. Bailey RG, Turner RD, Mullin N, Clarke N, Foster SJ, Hobbs JK. Article the Interplay between Cell Wall Mechanical Properties and the Cell Cycle in Staphylococcus aureus. Biophysical Society. 2014;107:2538–2545. Doi:10.1016/j.bpj.2014.10.036

46. Aungst BJ. Optimizing Oral Bioavailability in Drug Discovery: An Overview of Design and Testing Strategies and Formulation Options. Journal of Pharmaceutical Sciences. 2017;106(4):921–929. doi:10.1016/j.xphs.2016.12.002

47. Singh Y, Meher JG, Raval K, Khan FA, Chaurasia M, Jain NK, Chourasia MK. Nanoemulsion: Concepts, development and applications in drug delivery. Journal of Controlled Release. 2017;252():28–49. doi:10.1016/j.jconrel.2017.03.008

48. Lowy FD. Antimicrobial resistance: the example of Staphylococcus aureus. The Journal of Clinical Investigation. 2003;111(9):1265–1273. doi:10.1172/JCI18535

49. Audah KA, Ettin J, Darmadi J, Azizah NN, Anisa AS, Hermawan TDF, Tjampakasari CR, Heryanto R, Ismail IS, Batubara I. Indonesian Mangrove Sonneratia caseolaris Leaves Ethanol Extract Is a Potential Super Antioxidant and Anti Methicillin-Resistant Staphylococus aureus Drug. 2022;27(33):1–18. doi: 10.3390/molecules27238369.

